# MONI: A Pangenomics Index for Finding MEMs

**DOI:** 10.1101/2021.07.06.451246

**Authors:** Massimiliano Rossi, Marco Oliva, Ben Langmead, Travis Gagie, Christina Boucher

## Abstract

Recently, Gagie et al. proposed a version of the FM-index, called the *r*-index, that can store thousands of human genomes on a commodity computer. Then Kuhnle et al. showed how to build the *r*-index efficiently via a technique called prefix-free parsing (PFP) and demonstrated its effectiveness for exact pattern matching. Exact pattern matching can be leveraged to support approximate pattern matching but the *r*-index itself cannot support efficiently popular and important queries such as finding maximal exact matches (MEMs). To address this shortcoming, Bannai et al. introduced the concept of thresholds, and showed that storing them together with the *r*-index enables efficient MEM finding — but they did not say how to find those thresholds. We present a novel algorithm that applies PFP to build the *r*-index and find the thresholds simultaneously and in linear time and space with respect to the size of the prefix-free parse. Our implementation called MONI can rapidly find MEMs between reads and large sequence collections of highly repetitive sequences. Compared to other read aligners – PuffAligner, Bowtie2, BWA-MEM, and CHIC – MONI used 2–11 times less memory and was 2–32 times faster for index construction. Moreover, MONI was less than one thousandth the size of competing indexes for large collections of human chromosomes. Thus, MONI represents a major advance in our ability to perform MEM finding against very large collections of related references.

**Availability:** MONI is publicly available at https://github.com/maxrossi91/moni.

## 1 Introduction

In the past couple of decades, the cost of genome sequencing has decreased at an amazing rate, resulting in more ambitious sequencing projects including the 100K Genome Project (Turnbull et al., 2018) and the Vertebrate Genome Project (Rhie et al., 2021). Sequence read aligners – such as BWA (Li and Durbin, 2009), Bowtie (Langmead et al., 2009) and SOAP2 (Li et al., 2010) – have been fundamental methods for compiling and analyzing these and other datasets. Traditional read aligners build an index from a small number of reference genomes, find short exact matches between each read and the reference genome(s), and then extend these to find approximate matches for each entire read. Maximal exact matches (MEMs), which are exact matches between a read *R* and genome *G* that cannot be extended to the left or right^4^, have been shown to be the most effective seeds for alignment of both short reads (Li, 2013) and long reads (Miclotte et al., 2016; Vyverman et al., 2015). Hence, any index used for unbiased alignment should efficiently support finding these maximal exact matches (MEMs) and scale to indexing large numbers of genomes.

In recent years, we have come close to realizing such an index but some gaps still remain. The FM-index consists of the *Burrows-Wheeler transform* (BWT) of the input text, a rank data structure over that BWT and the suffix array (SA) sampled at regular intervals. Mäkinen and Navarro (2007) showed how to store the BWT and rank data structure in space proportional to the number *r* of runs in the BWT, which tends to grow very slowly as we add genomes to a genomic database, and still quickly count how many times patterns occur in the text. Because the product of the size of the SA sample and the time to locate each of those occurrences is at least linear in the size of the input text, however, Mäakinen and Navarro’s index is not a practical solution for alignment.

A decade later Gagie et al. (2020a) showed how to sample 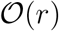 entries of the SA such that locating queries are fast. The combination of their SA sample with Mäkinen and Navarro’s rank data structure is called the *r*-index. However, Gagie et al. did not describe how to build the index – this was shown in a series of papers (Kuhnle et al., 2020; Mun et al., 2020; Boucher et al., 2019), which use *prefix-free parsing* (PFP) to preprocess the data in such a way that allows for the BWT and SA samples to be computed from the compressed representation. Exact pattern matching can be leveraged to support approximate pattern matching, by dividing patterns into pieces and searching for them separately, but the *r*-index itself cannot find MEMs. We cannot augment the *r*-index with the same auxiliary data structures that allow standard FM-indexes to find MEMs because, again, the product of their size and query time grows linearly with the text.

To address this shortcoming of the *r*-index, Bannai et al. (2020) describe a variant of the *r*-index that supports MEM-finding. More specifically, it finds the matching statistics of a pattern with respect to the indexed text, from which we can easily compute the MEMs, using a two-pass process: first, working right to left, for each suffix of the query string it finds a suffix of the text that matches for as long as possible; then, working left to right, it uses random access to the text to determine the length of those matches. We note that this computation of the matching statistics is enabled through the addition of a data structure that they refer to as *thresholds*. However, Bannai et al. did not say how to find those thresholds efficiently and until we have a practical construction algorithm we cannot say we really have a pangenomic index for MEM-finding.

In this paper, we show how to use PFP to find the thresholds at the same time as we build the *r*-index. We refer to the final data structure as MONI, from the Finnish for “multi”, since our ultimate intention is to index and use multiple genomes as a reference, whereas other approaches to pangenomics (Garrison et al., 2018; Li et al., 2020; Maarala et al., 2020) index models of genomic databases but not the databases themselves. We compare MONI to PuffAligner (Almodaresi et al., 2021), Bowtie2 (Langmead and Salzberg, 2012), BWA-MEM (Li, 2013), and CHIC (Valenzuela and Makinen, 2017) using GRCh37 and haplotypes taken from The 1000 Genomes Project Consortium (2015), and the *Salmonella* genomes taken from GenomeTrakr (Stevens et al., 2017). We show PuffAligner is between 1.7 and 4 slower for construction and uses between 3 and 12 times more memory for construction of the index for 32 or more haplotypes of chromosome 19. Bowtie2 is between 7 and 20 times slower for construction and uses between 2 and 15 times more memory for construction for these same datasets. BWA-MEM uses less memory but more time than Bowtie2 for construction. Only MONI and PuffAligner were capable of constructing an index for 1,000 haplotypes of chromosome 19 in less than 24 hours; BWA-MEM and Bowtie2 surpassed this. Moreover, MONI used 21GB of memory and 1.3 hours for construction of this index; whereas, PuffAligner used over 260 GB of memory and 5.2 hours for construction. Finally, the size of the data structure of PuffAligner was 1,114 times larger than that of MONI. Similarly, we show that MONI required significantly less time for construction than Bowtie2 and BWA-MEM, and also used significantly less memory than PuffAligner for construction of indexes of 100 or more *Salmonella* genomes. Finally, we demonstrate the use of indexing a larger number of genomes by comparing MEM-finding with a single reference genome, to that of 200 more genomes and show that we find MEMs (of length at least 75) for 4.6% more sequence reads.

We compare MONI to PuffAligner (Almodaresi et al., 2021), Bowtie2 (Langmead and Salzberg, 2012), BWA-MEM (Li, 2013), and VG (Garrison et al., 2018) using GRCh37 and from 10 to 200 complete genome haplotypes also taken from The 1000 Genomes Project Consortium (2015). We show that MONI and VG were the only tools able to build the index for 200 human genomes, in less than 48 hours, and using up to 756 GB of memory. In terms of wall clock time PuffAligner is the fastest to index 10 and 20 genomes, MONI is the fastest to index 50 genomes, and VG is the fastest to index 100 and 200 genomes. In terms of CPU time, MONI is always the fastest with a speedup that ranges from 5.92 to 1.16 with respect to the second fastest method. The peak memory usage of MONI is from 2.93 to 4.41 larger than the best method. The final data structure size of MONI is at most 1.18x larger than the final data structure of VG, and from 2.2 to 12.35 times smaller than the third smallest method. We aligned 611,400,000 100-bp reads from Mallick et al. (2016) against the successfully built indexes and evaluated the query time. PuffAligner is always the fastest, followed by Bowtie2, BWA-MEM, MONI and VG, while VG and MONI were the methods using the smallest amount of memory.

The rest of this paper is laid out as follows: in Section 2 we introduce some basic definitions, review prefix-free parsing, the definition of thresholds, and the computation of the matching statistics; in Section 3 we show how to compute the thresholds from the prefix-free parsing; in Section 4 we present our experimental results; and in Section 5 we discuss future work. For the sake of brevity we assume readers are familiar with SAs, BWTs, wavelet trees, FM-indexes, etc., and their use in bioinformatics. We refer the interested reader to Makinen et al. (2015) and Navarro (2016) for an in depth discussion on these topics.

## 2 Preliminaries

### 2.1 Basic Definitions

A *string S* = *S*[1..*n*] = *S*[1]*S*[2] ⋯ *S*[*n*] of length |*S*| = *n* is a sequence of *n* characters drawn from an alphabet *Σ* of size *σ*. We denote the empty string as *ε*. Given two indices 1 ≤ *i, j* ≤ *n*, we use *S*[*i..j*] to refer to the string *S*[*i*] ⋯ *S*[*j*] if *i* ≤ *j*, while *S*[*i..j*] = *ε* if *i* > *j*. We refer to *T* = *S*[*i..j*] as a *substring* of *S*, refer to *S*[1..*j*] as the *j*-th *prefix* of *S*, and refer to *S*[*i*..*n*] = *S*[*i*..] as the *i*-th *suffix* of *S*. A substring *T* of S is called *proper* if *T* ≠ *S*. Hence, a *proper suffix* is a suffix that is not equal to the whole string, and a *proper prefix* is a prefix that is not equal to the whole string.

### 2.2 SA, ISA and LCP

Given a text *S*, the *suffix array* (Manber and Myers, 1993) of *S*, denoted by SA*_S_*[1..*n*], is the permutation of {1, …, *n*} such that *S*[SA*_S_*[*i*]..*n*] is the *i*-th lexicographically smaller suffix of *S*. The *inverse suffix array* ISA*_S_*[1..*n*] is the inverse permutation of the suffix array, i.e. SA*_S_*[ISA*_S_*[*i*]] = *i*.

We denote the length of the longest common prefix (LCP) of *S* and *T* as `lcp`(*S, T*). And we define LCP array of *S* as LCP*_S_*[1..*n*] the array storing the values of the longest common prefix between two lexicographically consecutive suffixes of *S*, i.e. LCP*_S_*[1] = 0 and LCP*_S_*[*i*] = `lcp`(*S*[SA*_S_*[*i* – 1]..*n*], *S*[SA*_S_*[*i*]..*n*]) for all *i* = 2, …, *n*.

### 2.3 BWT, RLBWT and LF-mapping

The BWT (Burrows and Wheeler, 1994) of the text *S*, denoted by BWT*_S_*[1..*n*], is a reversible permutation of the characters of *S*. It is the last column of the matrix of the sorted rotations of the text *S*, and can be computed from the suffix array of *S* as BWT*_S_*[*i*] = *S*[SA*_S_*[*i*] – 1], where *S* is considered to be cyclic, i.e. *S*[0] = *S*[*n*]. The *LF-mapping* is a permutation on [1, *n*] such that SA*_S_*[LF(*i*)] = (SA*_S_*[*i*] – 1) mod *n*.

We represent the BWT*_S_*[1..*n*] with its run-length encoded representation RLBWT*_S_*[1..*r*], where *r* is the number of equal character runs of maximal size in the BWT*_S_*, e.g., runs of `A`’s, `C`’s and so forth. We write RLBWT*_S_*[*i*].*head* for the character of the *i*-th run of the BWT*_S_*, and RLBWT*_S_*[*i*].ℓ for its length.

When clear from the context, we remove the reference to the text *S* from the data structures, e.g., we write BWT instead of BWT*_S_* and SA instead of SA*_S_*.

### 2.4 Matching Statistics and Thresholds

The *matching statistics* of a string *R* with respect to *S* are an array of pairs saying, for each position in *R*, the length of the longest substring starting at that position that occurs in *S*, and the starting position in *S* of one of its occurrences. They are useful in a variety of bioinformatics tasks (Mäkinen et al., 2015), including computing the MEMs of *R* with respect to *S*.

#### Definition 1.

*The matching statistics of R with respect to S are an array M*[1..|*R*|] *of* (pos,len) *pairs such that: (1) S*[*M*[*i*].pos..*M*[*i*].pos + *M*[*i*].len – 1] = *R*[*i*..*i* + *M*[*i*].len – 1]; *and (2) R*[*i*..*i* + *M*[*i*].len] *does not occur in S*.

Suppose we have already computed *M*[*i* + 1].pos and now want to compute *M*[*i*].pos. Furthermore, suppose we know the position *q* in the BWT of *S*[*M*[*i* + 1].pos – 1] (or, equivalently, the position of *M*[*i* + 1].pos in SA). If BWT[*q*] = *R*[*i*], then we can set *M*[*i*].pos = *M*[*i* + 1].pos – 1, and the position in the BWT of *S*[*M*[*i*].pos – 1] is LF(*q*).

Bannai et al. (2020) observed that, if BWT[*q*] ≠ *R*[*i*], then we can choose as *M*[*i*].pos the position in *S* of either the last copy BWT[*q*′] of *R*[*i*] that precedes BWT[*q*], or the first copy BWT[*q*″] of *R*[*i*] that follows BWT[*q*], depending on which of the suffixes of *S* following those copies of *R*[*i*] has a longer common prefix with *S*[*M*[*i* + 1].pos..*n*]. For simplicity, we ignore here cases in which *q*′ or *q*″ are undefined.

They also pointed out that, if we consider BWT[*q*] moving from immediately after BWT[*q*′] to immediately before BWT[*q*″], then the length of the longest common prefix of the suffix of *S* following BWT[*q*] and the suffix of *S* following BWT[*q*′], is non-increasing; the length of the longest common prefix of the suffix of *S* following BWT[*q*] and the suffix of *S* following BWT[*q*″], is non-decreasing. Therefore, we can choose a threshold in that interval between BWT[*q*′] and BWT[*q*″] – that is, between two consecutive runs of copies of *R*[*i*] – such that if BWT[*q*] is above that threshold, then we can choose *M*[*i*].pos as the position in *S* of BWT[*q*′], and we can otherwise choose *M*[*i*].pos as the position in *S* of BWT[*q*″].

Bannai et al. proposed storing a rank data structure over the RL BWT of *S* so we can compute LF, the SA entry at the beginning and ending of each run in the BWT and, for each consecutive pair of runs of the same character, the position of a threshold in the interval between them. With these, we can compute *M*[1].pos, …, *M*[|*R*|].pos from right to left. We note they only said the thresholds exist and did not say how to find them efficiently; consequently, they did not give an implementation. They also proposed storing a data structure supporting random access to *S* so that, once we have all the pos values, we can compute the len values, from left to right: if we already have *M*[1].len, …, *M*[*i* – 1].len and now we want to compute *M*[*i* + 1].len; since *M*[*i*].len ≥ *M*[*i* – 1].len – 1, we can find *M*[*i*].len by comparing *S*[*M*[*i*].pos + *M*[*i* – 1].len – 1..|*S*|] to *R*[*i* + *M*[*i* – 1].len – 1..|*R*|] character by character until we find a mismatch. The number of characters we compare is a telescoping sum over i, so we use 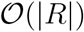 random accesses in total. Figure 1 shows an example of the computation of the matching statistics using the algorithm of Bannai et al.

**Fig. 1:**
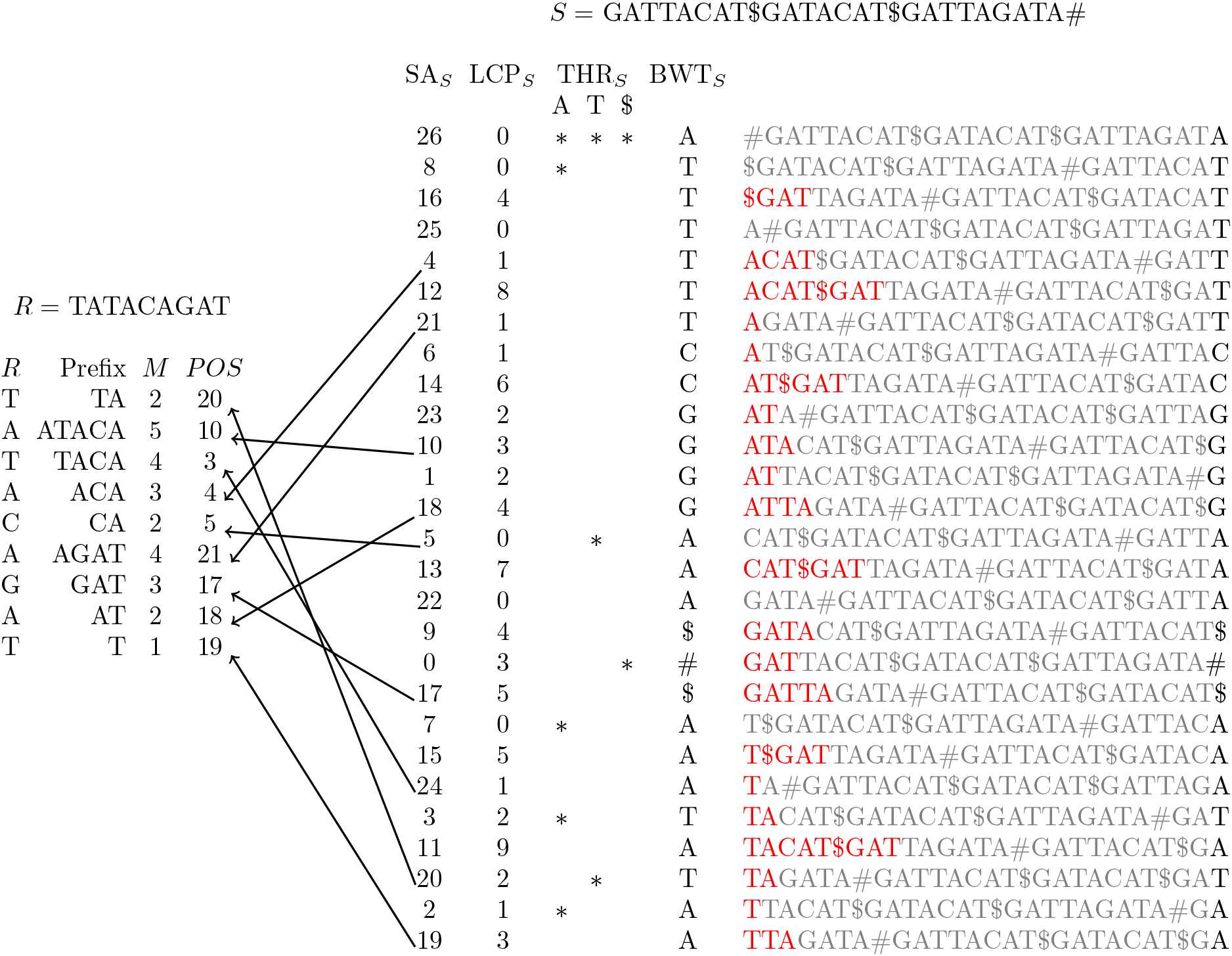
An illustration of the thresholds and matching statistics for identifying pattern *R* in the string *S*. Shown on the left is pattern *R*, the longest *Prefix* of the suffix of the pattern that occurs in *S*, its length in the matching statistics *M*, and the position of Prefix in the text. Shown on the right, continuing from left to right, is the SA*_S_*, LCP*_S_*, the thresholds THR*_S_* for the characters `A, T`, and `$`, the BWT*_S_*, and all rotations of *S*, with the longest common prefix between each consecutive rotations highlighted in red. We note that in practice we build the RLBWT and the SA*_S_* entries at the beginning and end of each run in the BWT*_S_*. For example, we would store {8, 21} for the first run of `T`’s in the BWT*_S_*. Moreover, the LCP*_S_* is just shown in red for illustrative purposes. The arrows illustrate the position in the *SA_S_* which each prefix corresponds to.

Examining Bannai et al.’s observations, we realized that we can choose the threshold between a consecutive pair of runs to be the position of the minimum LCP value in the interval between them. This allows us to build Bannai et al.’s index efficiently.

### 2.5 Prefix-free Parsing

Kuhnle et al. (2020) showed how to compute the *r*-index (i.e., RLBWT*_S_* and the SA entries at the starting and ending positions of runs in RLBWT*_S_*) using prefix-free parsing (PFP). For PFP, we parse *S* into overlapping phrases by passing a sliding window of length *w* over it and inserting a phrase boundary whenever the Karp-Rabin hashing of the contents of the window are 0 modulo a parameter. This can be done efficiently using only sequential access to *S*, so it works well in external memory, and it can also be parallelized easily. We call the substring contained in the window when we insert a phrase boundary a *trigger string* and we include it as both a suffix of the preceding phrase and a prefix of the next phrase. We treat *S* as cyclic, so the contents of the sliding window during the last w steps of the parsing are *S*[|*S*| – *w*..|*S*|], *S*[|*S*| – *w* + 1..|*S*|]*S*[1], …, *S*[|*S*|]*S*[1..*w* – 2]. It follows that: (a) each phrase starts with a trigger string, ends with a trigger string, and contains no other trigger strings; and (b) each character of *S* appears in exactly one phrase where it is not in the last *w* characters of that phrase.

The first property means no phrase suffix of length at least *w* is a proper prefix of any phrase suffix, which is the reason for PFP’s name. This and the second property mean that, for any pair of characters *S*[*i*] and *S*[*j*], we can compare lexicographically the suffixes starting at *S*[*i* + 1] and *S*[*j* + 1], again viewing *S* as cyclic, by (1) finding the unique phrases containing *S*[*i*] and *S*[*j*] not in their last *w* characters, (2) comparing the suffixes of those phrases starting at *S*[*i* + 1] and *S*[*j* + 1]; and (3) if those phrase suffixes are equal, comparing the suffixes of *S* starting at the next phrase boundaries.

#### Example 1.

Let *S* = GATTACAT#GATACAT#GATTAGATA and *w* = 2. Let *E* = {`AC, AG, T`#} be the set of strings where the Karp-Rabin hash is 0 modulo a parameter, therefore the *parsing* is *P* = *D*[1], *D*[2], *D*[4], *D*[2], *D*[5], *D*[3] and the *dictionary* is *D* = {##GATTAC, ACAT#, AGATA##, T#GATAC, T#GATTAG}. We note that in this example, the set of proper phrase suffixes of *D*[2] is {`AT#, T#, CAT#, #`}, and `AT#` and `CAT#` are the only one of length at least *w*.

#### Data Structures on Parse and Dictionary

We let *D* be the dictionary of distinct phrases and let *P* = *P*[1] ⋯ *P*[|*P*|] be the parse, viewed as a string of phrase identifiers. We define 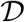 to be the text obtained by the concatenation of the phrases of the dictionary, i.e., 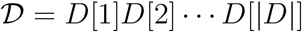. With a slight abuse of the notation, we refer to 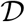 as *D* when it is clear from the context. We build the suffix array SA*_D_* of 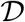, the inverse suffix array ISA*_D_* of 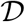, the longest common prefix array LCP*_D_* of 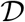, and a succinct range minimum query (RMQ) data structure on top of LCP*_D_*. Our last data structure we compute from *D* is a bitvector *b*_D_**[1..|*D*|] of length |*D*| which contains a 1 at the positions in 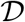 that correspond to the beginning of a phrase. We also provide *b*_D_** with rank and select support, i.e., rank*_q_*(*i*) and select*_q_*(*i*) returns the number of elements equal to *q* up to position *i* and select returns the position of the *i*-th occurrence of *q* where *q* is 0 or 1.

Finally, we build the suffix array SA*_P_* of *P*, the inverse suffix array ISA*_P_* of *P*, and the LCP*_P_* array of *P*. This leads to the following result describing the total time and space needed for the construction from *P* and *D*. Kuhnle et al. (2020) showed that the construction of BWT*_S_* and SA*_S_* from the dictionary and parse is linear in the size of the parse and dictionary. Moreover, the construction of the remaining structures is also linear (Navarro, 2016). Therefore, it follows that this data structure can be constructed in 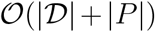 space and time since each individual structure can be constructed in time and space linear to the size of 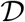 or *P*.

#### Computing the BWT

Suppose we lexicographically sort the distinct proper phrase suffixes of length at least *w*, and store the frequency of each such suffix in *S*. For each such phrase suffix *α*, all the characters preceding occurrences of *α* in *S* occur together in BWT*_S_*, and the starting position of the interval containing them is the total frequency in *S* of all such phrases suffixes lexicographically smaller than *α*. It may be that *α* is preceded by different characters in *S*, because *α* is a suffix of more than one distinct phrase, but then those characters’ order in BWT*_S_* is the same as the order of the phrases containing them in the BWT of *P*. These observations allow us to compute BWT*_S_* from *D* and *P* without decompressing them. Kuhnle et al. (2020) showed computing BWT*_S_* using PFP as a pre-processing step takes much less time and memory than computing it from *S* directly since *D* and *P* are significantly much smaller than *S*. The pseudocode for the construction algorithm of Kuhnle et al. is shown in Algorithm 1 in the Supplement.

#### Random Access to the SA

Now suppose, as shown by Boucher et al. (2021), we store a wavelet tree over the BWT of *P*, with the leaves labelled from left to right with the phrase identifiers in the co-lexicographic order of the phrases. For any phrase suffix *α*, all the identifiers of phrases ending with *α* appear consecutively in the wavelet tree; therefore, given an integer *j* and *α*, with the wavelet tree we can perform a 3-side range selection query to find the *j*th phrase in the BWT of *P* that ends with *α*. With some other auxiliary data structures whose sizes are proportional to the sizes of *D* and *P*, this lets us support random access to SA. We emphasize that we keep the wavelet tree and auxiliary data structures only during the construction of the *r*-index or Bannai et al.’s index, and that once we discard them we lose random access to SA. If we want to find SA[*i*], we first determine i) which such phrase suffix *α* follows BWT[*i*] in *S*, and ii) the lexicographic rank *j* of the suffix of *S* preceded by BWT[*i*], among all the suffixes of *S* prefixed by *α*, from the cumulative frequencies of the proper phrase suffixes of length at least *w*. We then use the wavelet tree to find the *j*th phrase in the BWT of *P* that ends with *α*. Finally, we use the suffix array of *P* to find that phrase’s position in *S* and compute the position of BWT[*i*] in *S*.

## 3 Computing Thresholds, Computing Matching Statistics and Finding MEMs

In the previous section we reviewed prefix free parsing. In this section we augment the prefix-free parsing based algorithm to compute the BWT and the SA samples to additionally compute the thresholds. Our description starts with our observation that the thresholds positions correspond to the position of a minimum of the LCP array in the interval between the two consecutive runs of the same character. This is our refinement of the definition. Next, we describe how to retrieve the thresholds. We show how to find MEMs from the matching statistics computed using Bannai et al.’s algorithm. Finally, we show implementation details on the threshold construction algorithm.

### 3.1 Redefining Thresholds

Here, we show that Bannai et al.’s thresholds are equivalent to the positions of a minimum LCP value in the interval between two consecutive runs of the same character. Given two suffixes *p*_1_ and *p*_2_ of *S*, we let *q*_1_ and *q*_2_ be their positions in the suffix array, i.e., *p*_1_ = SA*_S_*[*q*_1_] and *p*_2_ = SA*_S_*[*q*_2_]. We recall that the length of the longest common prefix between *S*[*p*_1_..*n*] and *S*[*p*_2_..*n*] can be computed as the minimum value of the LCP array of *S* in the interval LCP*_S_*[*q*_1_ + 1..*q*_2_], assuming w.l.o.g. that *q*_1_ < *q*_2_. Let MINLCP*_S_*[*q*_1_ + 1..q_2_] = min{LCP*_S_*[*q*_1_ + 1], …, LCP*_S_*[*q*_2_]}. This insight allows us to rewrite the definition of threshold in terms of LCP values as follows. Given a text *S*, let BWT*_S_*[*j*′..*j*] and BWT*_S_*[*k*..*k*′] be two consecutive runs of the same character in BWT*_S_*. From the definition of thresholds, we want to find a position *i* between positions *j* and *k*, such that for all *j* < *i*′ ≤ *i*, MINLCP*_S_*[*j* + 1..*i*′] is larger than or equal to MINLCP*_S_*[*i*′ + 1..*k*], while for all *i* < *i*′ ≤ *k*, MINLCP*_S_*[*i*′ + 1..*k*] is larger than or equal to MINLCP*_S_*[*j* + 1..*i*′]. This shows that position *i* is a threshold if the following holds:

– MINLCP*_S_*[*j* + 1..*i*] ≥ MINLCP*_S_*[*i* + 1..*k*], and
– MINLCP*_S_* [*j* + 1..*i* + 1] ≤ MINLCP*_S_*[*i* + 2..*k*],

assuming that MIN LCP*_S_*[*i* + 2..*k*] = ∞ if *i* > *k* – 2, i.e., *i* is the position of a minimum value in LCP*_S_*[*j* + 1..*k*]. This can be summarized by the following observation.

#### Observation 1

*Given text S, let* BWT*_S_*[*j*′..*j*] *and* BWT*_S_*[*k*..*k*′] *be two consecutive runs of the same character in* BWT*_S_*. *A position j* < *i* ≤ *k is a threshold if it corresponds to the minimum value in* LCP*_S_*[*j* + 1..*k*].

### 3.2 Computing Thresholds

We can find positions of minima in intervals in the LCP of *S* similarly to how we can compute the suffix array samples and, thus, compute Bannai et al.’s thresholds. If we want to find the position of a minimum in LCP[*i* + 1..*j*], we first check if BWT[*i*] and BWT[*j*] are followed in *S* by the same proper phrase suffix of length at least *w*. If they are not, we can find the position of the minimum from the LCP array of the proper phrase suffixes of length at least w: since the suffixes of *S* following BWT[*i*] and BWT[*j*] are not prefixes of each other, their longest common prefix is a proper prefix of both of them. The situation is more complicated when BWT[*i*] and BWT[*j*] are followed in *S* by the same proper phrase suffix *α* of length at least *w*. First, let us consider the simpler problem of finding the length of the longest common prefix of the suffixes of *S* following BWT[*i*] and BWT[*j*], using some more auxiliary data structures. From SA[*i*] and SA[*j*] we can find the phrases containing BWT[*i*] and BWT[*j*] in *S*. Using the inverse suffix array of *P*, we find the lexicographic rank of the the suffixes of *S* starting at the next phrase boundaries after BWT[*i*] and BWT[*j*], among all suffixes of *S* starting at phrase boundaries. Using a range-minimum data structure over all |*P*| such suffixes of *S*, we find the length of the longest common prefix of those two suffixes. Finally, we add |*α*| – *w*, the length of the phrase suffixes after BWT[*i*] and BWT[*j*] minus the length of their overlaps with next phrases.

The range minimum query mentioned above gives us the position of a minimum in LCP[ISA[SA[*i*] + |*α*| – *w*] + 1..ISA[SA[*j*] + |*α*| – *w*]], which could be a much wider interval than LCP[*i* + 1..*j*]. To see why, consider that each of the suffixes of *S* starting at one of the positions SA[*i*], …, SA[*j*] consists of *α*[1..|*α*| – *w*] followed by a suffix starting at one of the positions SA[*i*] + |*α*| – *w*, …, SA[*j*] + |*α*| – *w*, but not all the suffixes starting at the latter positions are necessarily preceded by *α*[1..|*α*| – *w*]. We find the position of a minimum in LCP[*i* + 1..*j*] by filtering out the positions SA[*i*] + |*α*| – *w*, …, SA[*j*] + |*α*|– *w* in *S* that are not preceded by *α*[1..|*α*| – *w*]: we find the value *t* in [*i* + 1..*j*] such that LCP[ISA[SA[tb] + |*α*| – *w*]] is after the minimum in LCP[ISA[SA[*i*] + |*α*| – *w*] + 1..ISA[SA[*j*] + |*α*| – *w*]]. We can do this efficiently using the wavelet tree over the BWT of *P*: the position of a minimum in LCP[ISA[SA[*i*] + |*α*| – *w*] + 1..ISA[SA[*j*] + |*α*| – *w*]] corresponds to a certain phrase boundary in *S*, and thus to the identifier in the BWT of *P* for the phrase preceding that phrase boundary; we are looking for the next identifier in the BWT of *P* for a phrase ending with *α*, which we can find quickly because the phrase identifiers are assigned to the leaves of the wavelet tree in co-lexicographic order.

### 3.3 Computing Matching Statistics and Finding MEMs

Given a query string *R*, we compute the matching statistics of *R* from the thresholds using the algorithm of Bannai et al. given in Subsection 2.4. Next, it follows that we can find the MEMs for *R* by storing the non-decreasing values of the lengths of the matching statistics. This is summarized in the following lemma.

#### Lemma 1.

*Given input text S*[1..*n*] *and query string R*[1..*m*], *let M*[1..*m*] *be the matching statistics of R against S*. *For all* 1 < *i* ≤ *m, R*[*i*..*i* + ℓ – 1] *is a maximal exact match of length* ℓ *in S if and only if M*[*i*] = ℓ *and M* [*i* – 1] ≤ *M*[*i*].

*Proof*. First, we show that if *R*[*i*..*i* + ℓ – 1] is a maximal exact match of length ℓ in *S* then *M*[*i*] = ℓ and *M*[*i* – 1] ≤ *M*[*i*]. Since *R*[*i*..*i* + ℓ – 1] is a MEM then there exists a *j* such that *R*[*i*..*i* + *ℓ* – 1] = *S*[*j*..*j* + *ℓ* – 1] and hence, *M*[*i*] ≤ ℓ. Moreover, ℓ is the length of the longest prefix of *R*[*i*..*m*] that occurs in *S* since *R*[*i*..*i* + ℓ] does not occur in *S*, i.e., *M*[*i*] = ℓ. It also holds that *R*[*i* – 1..*i* + ℓ – 1] does not occur in *S*. This implies that the length of the longest common prefix of *R*[*i* – 1..*n*] that occurs in *S* is smaller than ℓ +1, and therefore, *M*[*i* – 1] ≤ ℓ = *M*[*i*].

Next, we prove the other direction. Given *M*[*i*] = ℓ, by definition of matching statistics, there exists a *j* such that *S*[*j*..*j* + ℓ – 1] = *R*[*i*..*i* + ℓ – 1] and *R*[*i*..*i* + ℓ] does not occur in *S*. We also have that *R*[*i* – 1..*i* – 1 + *M*[*i* – 1]] does not occur in *S*. Since *M*[*i* – 1] ≤ *M*[*i*] then also *R*[*i* – 1..*i* – 1 + *M*[*i*]], implying *R*[*i*..*i* + ℓ – 1] is a maximal exact match.

### 3.4 Implementation

We implemented MONI using the bitvectors of the `sdsl-lite` library (Gog et al., 2014) and their rank and select supports. We used SACA-K (Nong, 2013) to lexicographically sort the parses, and gSACA-K (Louza et al., 2017) to compute the SA and LCP array of the dictionary. We provide random access to the reference using the practical random access to SLPs of Gagie et al. (2020b) built on the grammar obtained using BigRePair (Gagie et al., 2019). We used the `ksw2` (Suzuki and Kasahara, 2018; Li, 2018) library available at https://github.com/lh3/ksw2 to compute the local alignment.

#### Removing the Wavelet Tree

In Section 2, for sake of explanation, we used a wavelet tree to provide random access to SA*_S_*, but, as shown by Kuhnle et al. (2020), to build BWT*_S_*, we only need sequential access to SA*_S_*. To provide sequential access to SA*_S_* we store an array called *inverted list* which stores a list for each phrase in *P* of the sorted occurrences of the phrases in BWT*_P_*. When processing the proper phrase suffixes in lexicographic order, if a proper phrase suffix *α* is suffix of more than one phrase, and is preceded by more than one character, we merge the lists of the occurrences of the phrases that contain *α* in the BWT of *P* using a heap. Since we want to find the phrases in the BWT of *P* ending with *α* in order, it is enough to scan the elements of the merged lists in increasing order.

The wavelet tree is also used to find the position of the minimum *LCP_S_* in a given interval, i.e., to perform a range minimum query. First we store a variation of the LCP array of *P*, we called SLCP, which we define as follows. We let SLCP[1] = 0. Next, for all 1 < *i* < |*P*|, we let SLCP[*i*] be equal to `lcp`(*S*[*p_i_*..*n*], *S*[*p*_i-1_..*n*]), where *p_i_* and *p*_i-1_ are the positions in SA*_S_* of the beginning of the lexicographically *i*-th and *i* – 1-th phrases of the parse. The SLCP array can be computed in 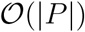-time with a slight modification of the algorithm of Kasai et al. (2001). Next, we build a succinct range minimum query (RMQ) data structure from the SLCP.

Given the SLCP array, the following lemma shows how to compute the LCP values,

##### Lemma 2.

*Given the prefix-free parsing P of S*[1..*n*] *with dictionary D, for all* 1 < *i* ≤ *n*, *let α and β be the unique proper phrase suffixes of length at least w of the phrases which* SA*_S_* [*i* – 1] *an* SA*_S_*[*i*] *belongs to, and let p*_1_ *and p*_2_ *be the positions of the phrases which α and β belongs to, in* BWT*_P_*. *Assuming w.l.o.g. that p*_1_ < *p*_2_, *then*

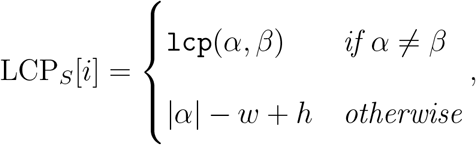

*where* h = min{SLCP[*p*_1_ + 1], …, SLCP[*p*_2_]}.

*Proof*. First, we consider the case where *α* ≠ *β*. Since 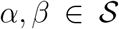, where 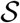 is the prefix-free set of proper phrase suffixes of length at least *w*, then LCP*_S_*[*i*] = `lcp`(*α, β*). In the second case, i.e. if *α* = *β*, LCP*_S_*[*i*] = |*α*| + `lcp`(*S*[SA*_S_*[*i* – 1] + |*α*|..*n*],S[SA*_S_*[*i*] + |*α*|..*n*]). We note that *S*[SA*_S_*[*i* – 1] + |*α*|..*n*] and *S*[SA*_S_*[*i*] + |*α*|..*n*] are suffixes of *S* starting at phrase boundaries, hence their longest common prefix can be computed using SLCP as follows. Let *h* = min{SLCP[*p*_1_ + 1], …, SLCP[*p*_2_]}. We have to show that `lcp`(*S*[SA*_S_*[*i* – 1] + |*α*|..*n*], *S*[SA*_S_*[*i*] + |*α*|..*n*]) = *h*. Since the phrases in *P* are represented by their lexicographic rank in *D*, the relative lexicographic rank of suffixes of *P* is the same as the relative lexico-graphic rank of their corresponding suffixes on *S*. Hence, the value of SLCP[*i*] corresponds the minimum value in the LCP interval between the positions in SA*_S_* of the suffixes starting with phrases SA*_P_*[*i* – 1] and SA*_P_*[*i*]. Thus, computing the minimum value in SLCP[*j*..*i*] is equivalent to computing the minimum value in the LCP interval between the positions in SA*_S_* of the suffixes starting with phrases SA*_P_*[*j*] and SA*_P_*[*i*]. Then LCP*_S_*[*i*] = |*α*| – *w* + *h*. We subtract *w* to the length because the last *w* character of *α* are the same as the first *w* characters of the phrase following *α*, that are included in the values of SLCP.

We observe that, during the construction of the thresholds, we do not need to answer arbitrary range minimum queries, but only those between two equal-letter runs. In addition we are building the BWT sequentially. Hence, while building the BWT, we can store, for each character of the alphabet, the position of the minimum LCP in the interval starting at the last occurrence of the character. We can clearly compute all the values of LCP*_S_*, while building the BWT, however this would require to visit, for each proper phrase suffix, all the occurrences of the phrases containing it, in the BWT of *P*. This can be avoided by noticing that if a proper phrase suffix is always preceded by the same character, the minimum LCP value in the interval is in position of the first suffix, because the previous suffix starts with a different proper phrase suffix. For the other LCP values, we use the positions of the phrases in the BWT of *P* that are computed using the inverted list, and we use the RMQ over the SLCP to compute the length of the longest common prefix between two consecutive suffixes of *S*. We also note that the *P*, SA*_P_* and ISA*_P_* are used only during the construction of SLCP and the inverted list and therefore, are discarded after this construction.

Algorithm 2 in the Supplement gives the pseudocode for building the thresholds.

## 4 Experiments

We demonstrate the performance of MONI through the following experiments: (a) comparison between our method and general data structures and algorithms that can calculate thresholds, (b) comparison between the size and time required to build the index of MONI and that of competing read aligners, and (c) comparison between MONI and competing read aligners with respect to their alignment performance.

The experiments were performed on a server with Intel(R) Xeon(R) CPU E5-2640 v4 @ 2.40GHz with 40 cores and 756 gigabytes of RAM running Ubuntu 16.04 (64bit, kernel 4.4.0). The compiler was `g++` version 5.4.0 with `-03 -DNDEBUG -funroll-loops -msse4.2` options. The running time was found using the C++11 `high_resolution_clock` facility and memory usage was found with the `malloc_count` tool (https://github.com/bingmann/malloc_count) when available, otherwise we used the maximum resident set size provided by `/usr/bin/time`. Where not specified, we refer to wall clock time as runtime. All experiments that either exceeded 24 hours or requires more than 756 GB of RAM were omitted from further consideration, e.g. `chr19.1,000` and `salmonella.10,000` for `gsacak` and `sdsl`.

### 4.1 Datasets

We used the following data for our experiments: *Salmonella* genomes taken from Genome-Trakr (Stevens et al., 2017), and sets of haplotypes from The 1000 Genomes Project Consortium (2015). In particular, we used collections of 50, 100, 500, 1,000, 5,000, and 10, 000 *Salmonella* genomes, where each collection is a superset of the previous. We denote these as `salmonella.50,..,salmonella.10,000`. We created a collection of chromosome 19 haplotypes using the `bcftools consensus` tool to integrate variant calls from the phase-3 callset into chromosome 19 of the GRCh37 reference. We did this for sets of 1, 2, 4, 8, 16, 32, 64, 128, 256, 512, and 1, 000 distinct haplotypes, where each set is a superset of the previous. We denote these as `chr19.1,..,chr19.1,000`. Lastly, we repeated this for the whole human genome and obtained sets of 1, 10, 20, 50, 100, and 200 distinct haplotypes, where each set is a superset of the previous. We denote these as `HG, HG.10, HG.20, HG.50, HG.100` and `HG.200`. All DNA characters in the reference besides A, C, G, T and N were removed from the sequences before construction.

### 4.2 Competing Read Aligners

We compare MONI to Bowtie2 (Langmead and Salzberg, 2012) (v2.4.2) and BWA-MEM (Li and Durbin, 2009) (v0.7.17), and to more recent tools that have demonstrated efficient alignment to repetitive text, i.e., PuffAligner (Almodaresi et al., 2021) (v1.0.0) and CHIC (Valenzuela et al., 2018) (v0.1). PuffAligner was released in 2020 and compared against deBGA (Liu et al., 2016), STAR (Dobin et al., 2013), and Bowtie2. We build Bowtie2, BWA-MEM, and Puffaligner using the default options, while we build CHIC using the relative Lempel-Ziv parsing method `--lz-parsing-method=RLZ` with 10% as prefix length of text from which phrases in the parse can be sourced, and we fixed the maximum pattern length to be 100. We tested CHIC with both BWA-MEM and Bowtie2 as kernel manager, which we denote as chic-bwa and chic-bowtie2. We run all methods using 32 threads, except the construction of the BWA-MEM index where multi-threading is not supported.

### 4.3 Comparison to General Data Structures: Thresholds Construction

We compare MONI to other data structures that can compute thresholds. First, we compute the thresholds using the minimum LCP value in each run of the BWT, which we build using the LCP construction algorithm of Prezza and Rosone (2019) that is available at https://github.com/nicolaprezza/rlbwt2lcp. We denote this method as `bwt2lcp`. Next, we compute the thresholds directly from the LCP array computed using gSACA-K (Louza et al., 2017). We denote this method as `gsacak`. Both methods take as input the text *S* and provide as output the BWT*_S_*, the samples of SA*_S_* at the beginning and at the end of a BWT run, and the thresholds. Hence, MONI includes the construction of the prefix-free parsing using the parsing algorithm of BigBWT (Boucher et al., 2019), while `bwt2lcp` includes the construction of the BWT and the samples using BigBWT. In both cases, BigBWT is executed with 32 threads, window size *w* = 10, and parameter *p* = 100. We ran each algorithm five times for sets of chromosome 19 up to 64 distinct haplotypes.

We compare MONI, `gsacak` and `bwt2lcp` with respect to the time and peak memory required to construct the thresholds using the Chromosome 19 dataset. Figure 2 illustrates these values. We can observe that MONI is fastest except for very small datasets, i.e., `chr19.1` and `chr19.2`, where `gsacak` is fastest. MONI’s highest speedup is 8.9x with respect to `bwt2lcp` on `chr19.1,000` and 17.3x with respect to `gsacak` on `chr19.512. bwt2lcp` uses the least memory for collections up to 32 sequences, but MONI uses the least for larger collections.

**Fig. 2:**
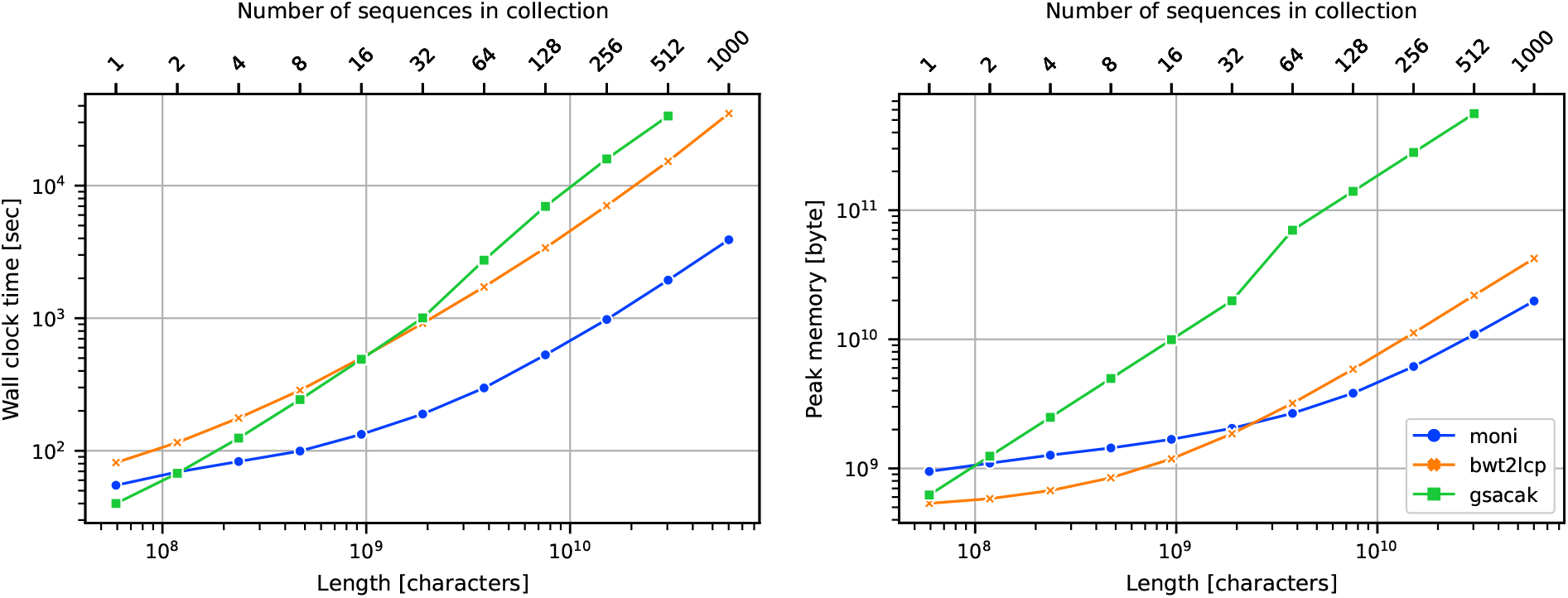
Chromosome 19 dataset thresholds construction running time (**left**) and peak memory (**right**).

On the *Salmonella* dataset, in Figure 3 we report that MONI is always the fastest with the highest speedup of 4.1x with respect to `gsacak` on `salmonella.5000` and 3.1x with respect to `bwt2lcp` on `salmonella.10,000`. On the other hand, `bwt2lcp` always uses less memory than MONI and `gsacak`. The high memory consumption of MONI on `salmonella` is mainly due to the size of the dictionary for `salmonella`, that is about 16 times larger than the dictionary for chromosome 19.

**Fig. 3:**
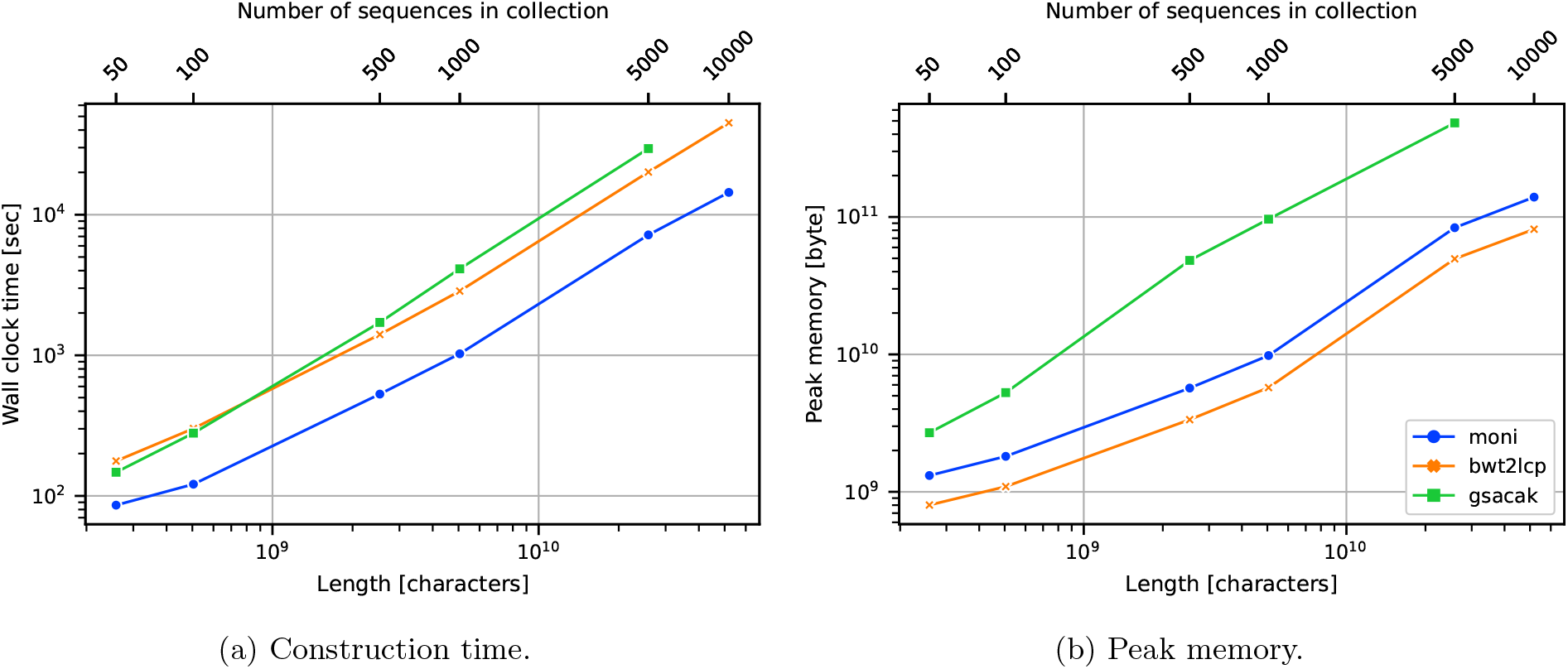
*Salmonella* dataset thresholds construction running time (**left**) and peak memory usage (**right**).

### 4.4 MEM-finding in a Pangenomics Index

Next, we evaluated the degree to which indexing more genomes allowed MONI to find longer MEMs and thus, more reliable anchors for alignment. We used MONI to build the thresholds for the GRCh37 reference genome (`HG`), and for GRCh37 and *i* – 1 randomly selected haplotypes from the 1000 Genomes Project phase-3 callset, which we denote as `HG.i` for *i* = 10, 20, 50, 100, 200. Next, we computed the MEMs for 611,400,000 100-bp reads from Mallick et al. (2016) (Accession number ERR1019034_1). Table 1 shows the number of reads having a MEM of length at least 25, 50, and 75. For larger collections, we also measured the number of additional reads having a MEM of each length with respect to the next-smaller collection (“+Reads”). For example, MONI was able to find over an additional 10,452,669 MEMs of length at least 75 when using the `HG.10` collection compared to using `HG`, an increase of 4.12%. This demonstrates the utility of indexing a set of reference genomes rather than a single genome: reads tend to have longer MEMs, corresponding to longer uninterrupted stretches of genetically identical sequence between the read and a closely related reference in the set.

**Table 1:** The number of reads containing a MEM of minimum length 25, 50, and 75 for the reference genome, 10, 20, 50, 100, and 200 haplotypes. We give the total number of reads that have a MEM (“No. of Reads”) and the number of additional reads that have a MEM (“+Reads”). We note that +Reads compares the number of MEMs of the current to the next-previous set, i.e., `HG.100 - HG.50` and so forth.

| MEM | HG | HG.10 |  | HG.20 |  | HG.50 |  | HG.100 |  | HG.200 |  |
| --- | --- | --- | --- | --- | --- | --- | --- | --- | --- | --- | --- |
|  | No. of Reads | No. of Reads | +Reads | No. of Reads | +Reads | No. of Reads | +Reads | No. of Reads | +Reads | No. of Reads | +Reads |
| 25 | 411,665,608 | 412,292,276 | 626,668 | 412,383,562 | 91,286 | 412,580,107 | 196,545 | 412,678,277 | 98,170 | 412,818,387 | 140,110 |
| 50 | 309,876,128 | 311,825,333 | 1,949,205 | 311,986,530 | 161,197 | 312,172,012 | 185,482 | 312,298,469 | 126,457 | 312,460,499 | 162,030 |
| 75 | 253,953,551 | 264,406,220 | 10,452,669 | 264,941,235 | 535,015 | 265,311,230 | 369,995 | 265,510,475 | 199,245 | 265,770,839 | 260,364 |

### 4.5 Comparison to Read Aligners: Construction Space and Time

We compare MONI to competing read aligners with respect to the time used for construction, peak memory used for construction, and size of the resulting data structure on disk. Figures 4 and 5 illustrate these comparisons for the human chromosome 19 and *Salmonella* datasets, respectively. For chromosome 19, MONI is faster than Bowtie2, BWA-MEM, and PuffAligner for 16 or more copies of chromosome 19. In particular for 16 or more copies of chromosome 19, MONI is between 1.6 and 4 times faster than PuffAligner, 3.8 and 32.8 times faster than BWA-MEM, and 4.6 and 20.6 times faster than Bowtie2. Bowtie2 and BWA-MEM were only faster than MONI on `chr19.1` and `chr19.2`. For small input (i.e., `chr19.1` to `chr19.8`), there was negligible difference between all the methods, i.e., less than 200 CPU seconds. Bowtie2 and BWA-MEM are not shown for `chr19.1000` in Figure 4 because they required over 24 hours for construction. MONI has lower peak memory usage compared to BWA-MEM for more than 32 copies of chromosome 19, to Bowtie2 for more than 8 copies of chromosome 19, and to PuffAligner when the number copies for chromosome 19 exceeded 8. Bowtie2 used between 1.2 and 14 times more memory than MONI, BWA-MEM used between 1.1 and 3.8 times more, and PuffAligner used between 1.7 and 12 times more. For small input (i.e., `chr19.1`, `chr19.2`, and `chr19.8`) there was negligible difference between the methods (i.e., less than 1GB). In addition, MONI’s data structure was the smallest for all experiments using chromosome 19. The index of Bowtie2, BWA-MEM and PuffAligner was between 2.8 and 945, 3.3 and 913, and between 9.8 and 1,114 times larger than ours. PuffAligner consistently had the largest index. Although chic-bwa and chic-bowtie had competitive construction time and produced smaller indexes compared to Bowtie2, BWA-MEM and PuffAligner, they required more memory and produced larger indexes compared to MONI. Moreover, the chic-based methods were unable to index more than 32 copies of chromosome 19; after this point it truncated the sequences.

**Fig. 4:**
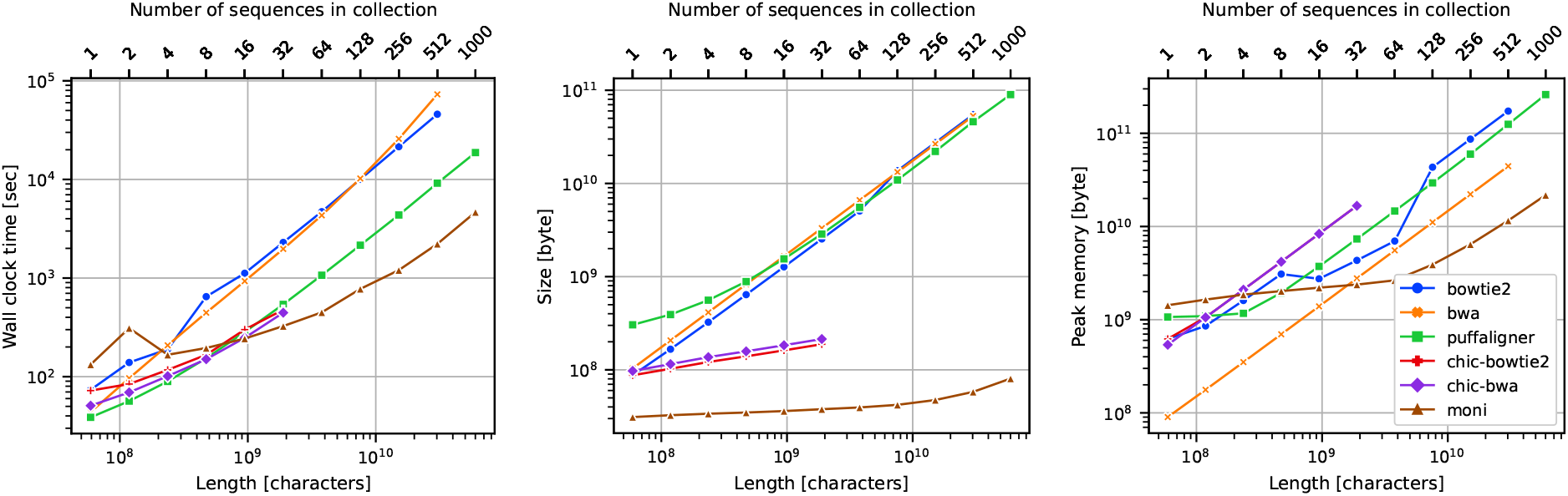
Chromosome 19 dataset index construction running time (**left**), index size (**center**), and peak memory usage (**right**).

**Fig. 5:**
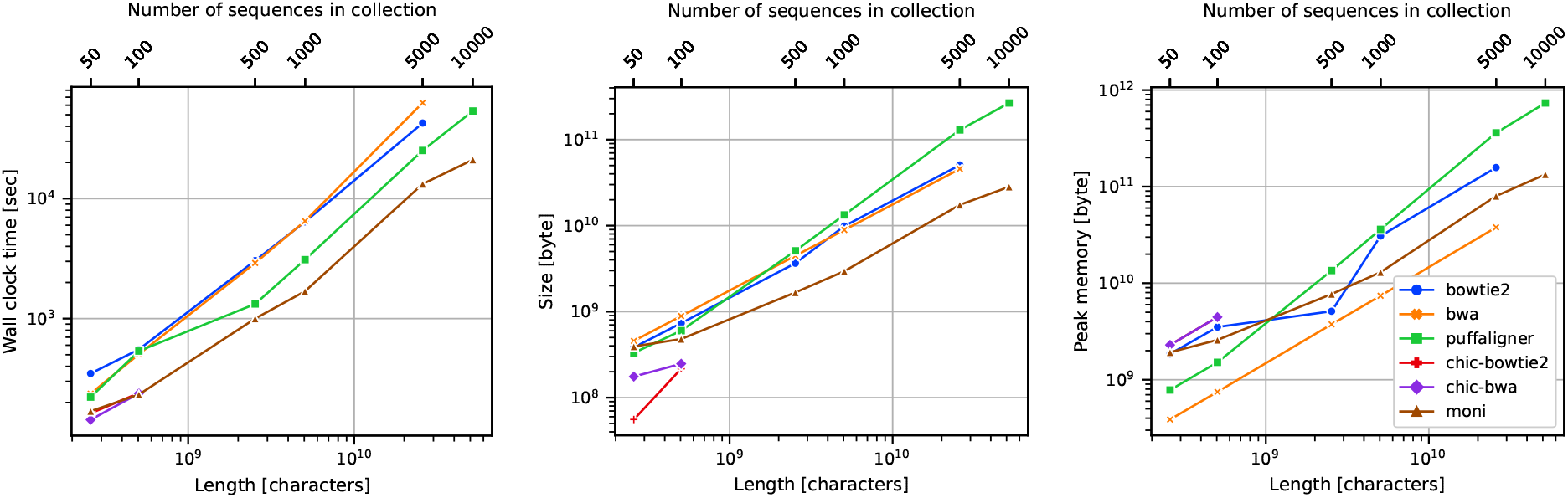
*Salmonella* dataset index construction running time (**left**), index size (**center**), and peak memory usage (**right**).

Results for *Salmonella* are similar to those for chromosome 19. MONI was faster and used less memory for construction compared to PuffAligner, Bowtie2, and BWA-MEM for all sets of salmonella greater than 50; the one exception is `salmonella.500`, where Bowtie2 used approximately 2 GB less memory for construction. Our final data structure had a consistently smaller disk footprint compared to indexes for BWA-MEM, Bowtie2, and PuffAligner for 100 or more strains of *Salmonella*. For `salmonella.50` the difference in size between MONI, PuffAligner, Bowtie2 and BWA-MEM was negligible. Although, chic-bwa and chic-bowtie were competitive with respect to construction time and size, they truncated the sequences after there was more than 100 strains of *Salmonella*. Bowtie2 and BWA-MEM are not shown for `salmonella.10000` in Figure 5 because they required over 24 hours for construction.

In summary, MONI was the most efficient with respect to construction time and memory usage for 32 or more copies of chromosome 19. In general, PuffAligner had faster construction time than Bowtie2 and BWA-MEM but had higher peak memory usage than BWA-MEM. PuffAligner and Bowtie2 had comparable peak memory usage. BWA-MEM had the most competitive peak memory usage to MONI but had the longest construction time for larger inputs, i.e. for 128 or more chromosome 19 haplotypes and 1,000 or more strains of *Salmonella*.

### 4.6 Comparison to Short Read Aligners: Human Pangenome

We attempted to index the `HG, HG.10, HG.20, HG.50, HG.100`, and `HG.200` collections using MONI, BWA-MEM, Bowtie2, PuffAligner, and VG. We recorded the wall clock time, CPU time, and the maximum resident set size using `/usr/bin/time`. All tools that used more than 48 hours of wall clock time or exceeded a peak memory footprint of 756 GB were omitted from further consideration. BWA-MEM required more than 48 hours of wall clock time to index `HG.20`, Bowtie2 required more than 48 hours of wall clock time to index `HG.50`, and PuffAligner required more than 756 GB of main memory to index `HG.200`.

BWA-MEM, Bowtie2, PuffAligner, and VG all report alignments in SAM format. Though MONI is not a full-featured aligner, we implemented a dynamic programming algorithm to extend MEMs as a fairer comparison. MONI joins MEMs as follows: (1) we compute matching statistics and MEMs for a given read using our index, (2) if the read has a MEM of length at least 25, we extract the MEM plus the 100-bp regions on either side from the corresponding reference, and lastly, (3) we perform Smith-Waterman alignment (Smith and Waterman, 1981) using a match score of 2, a mismatch penalty of 2, a gap open penalty of 5, and a gap extension penalty of 2. We consider aligned a read with a Smith-Waterman score greater than 20 + 8 log(*m*), where *m* is the length of the read.

Using the aligners and collections for which we could successfully build an index, we aligned the 611,400,000 reads from Mallick et al. (2016) and measured the wall clock time, the CPU time and the peak memory footprint required for alignment (Figure 6). All tools were run using 32 simultaneous threads. All methods successfully built indexes for `HG.10`. MONI was the fastest tool to build the index when considering the CPU time (7 hours), and the second fastest when considering the wall clock time (6 hours and 45 minutes), after PuffAligner. The tools produced indexes ranging in size from 24.06 to 58.91 GB. MONI’s index was the smallest. Peak memory footprint varied from 44.92 to 172.49 GB, with MONI having the fourth smallest (131.83 GB). On queries, BWA-MEM’s large footprint (128.22 GB) is likely due to the fact that it was run with 32 simultaneous threads. Wall clock times required ranged from more than 1 to less than 12 hours, with MONI being the second slowest (7 hours and 54 minutes). CPU time required ranged from more than 1 to more than 15 days with MONI being the the second slowest (7 days and 18 hours). For the `HG.100` collection, only MONI, PuffAligner, and VG were able to build an index, with VG being the fastest to build the index in terms of wall clock time (14 hours and 36 minutes), and MONI was the second fastest (20 hours and 15 minutes). MONI was the fastest when considering the CPU time (1 day and 2 hours). VG used the smallest amount of peak memory (90.65 GB) and final data structure size (25.93 GB). MONI used 3.08 times more peak memory (206.25 GB), but the final data structure size is only 3.8 GB larger (29.80 GB). PuffAligner used the largest amount of both peak memory (782.78 GB) and final data structure size (368.22 GB). On queries, PuffAligner is the fastest on both wall-clock (4 hours and 32 minutes) and CPU time (2 day and 23 hours) but used the largest amount of peak memory (329.18 GB), while MONI was the second fastest on both wall clock (8 hours and 31 minutes) and CPU time (8 days and 10 hours), and used the second smallest amount of memory (29.57 GB). VG was the slowest on both wall clock time (12 hours and 25 minutes) and CPU time (16 days and 14 hours), but used the smallest amount of peak memory (26.56 GB). For the `HG.200` collection, only MONI and VG were able to build an index, with VG being the fastest in terms of wall clock time (15 hours and 41 minutes), and MONI being the fastest in terms of CPU time (2 days and 6 hours). VG used the smallest amount of peak memory (94.65 GB) and had the smallest final data structure size (26.88 GB). On queries, MONI was the fastest (8 hours and 35 minutes), while VG used the smallest amount of peak memory (27.84 GB).

**Fig. 6:**
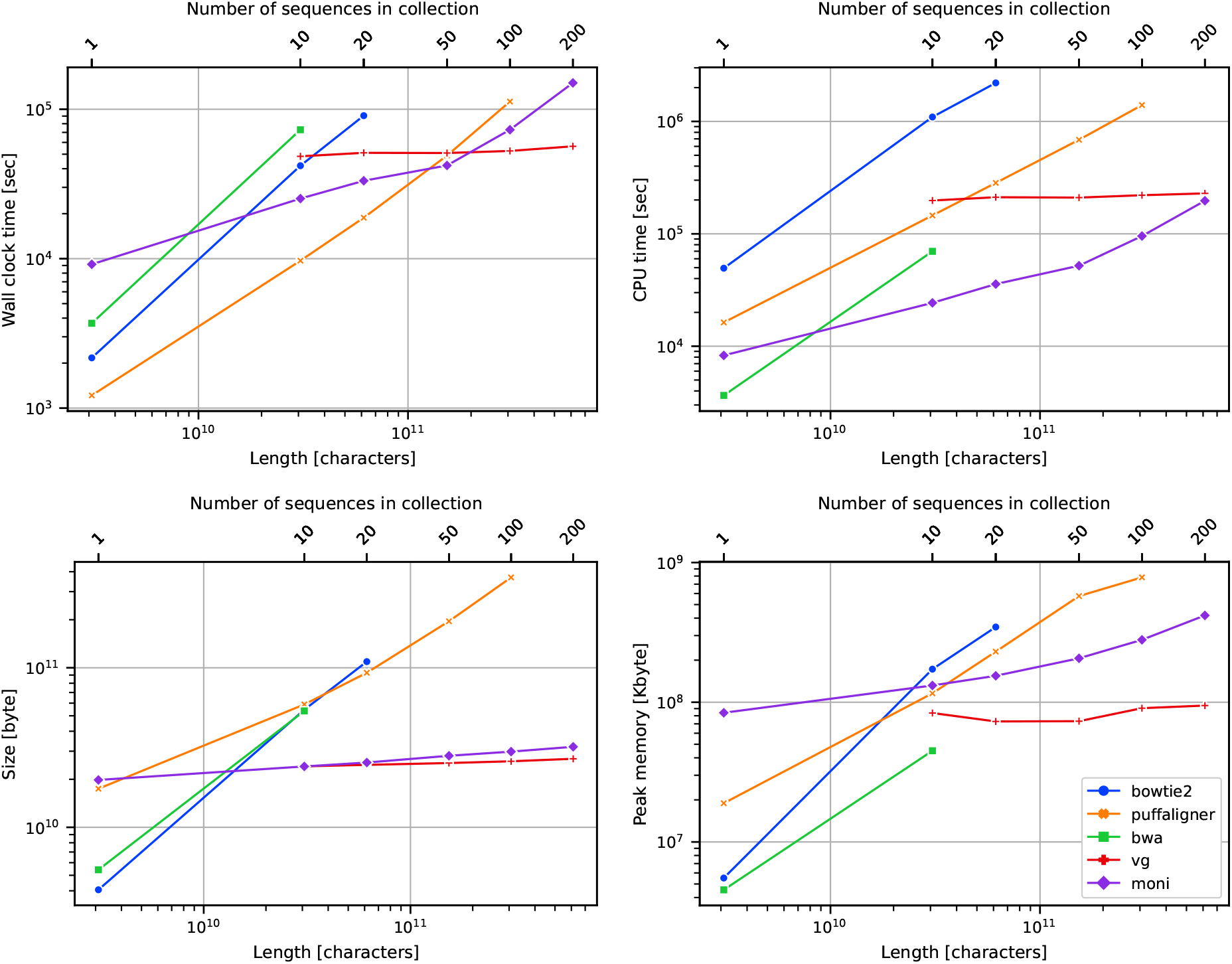
Human genomes dataset index construction wall clock time (**upper left**), CPU time (**upper right**), index size (**lower left**), and peak memory usage (**lower right**).

**Fig. 7:**
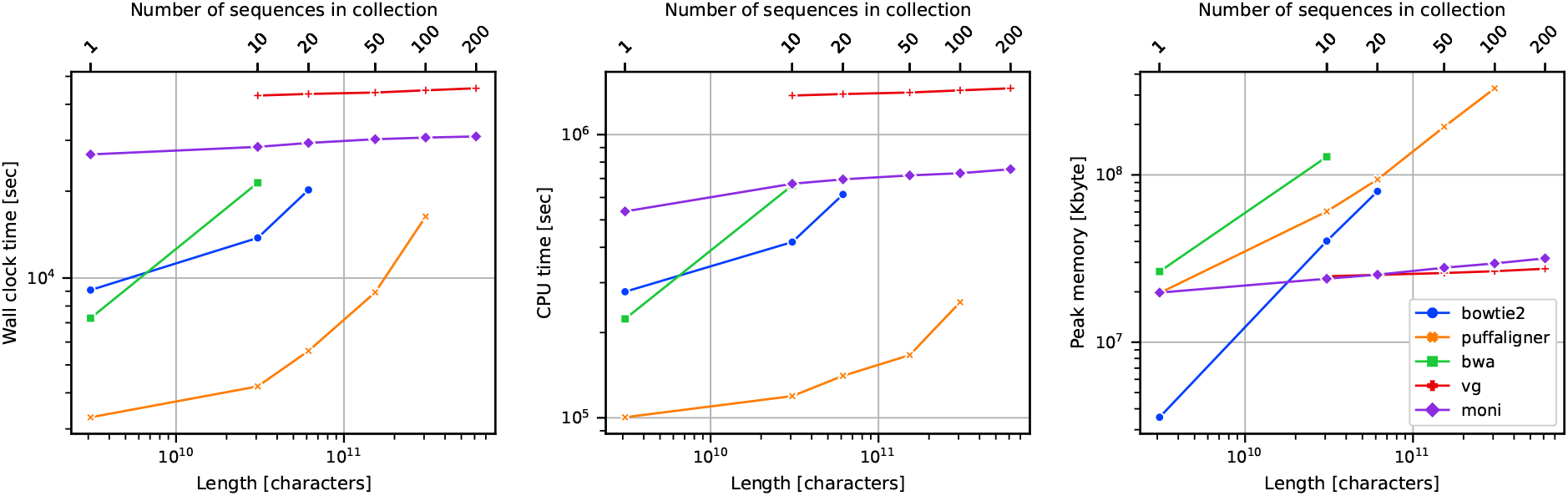
Human genomes query wall clock time (**left**), CPU time (**center**), and peak memory usage (**right**).

We note that MONI running time heavily depends on the total length of the input reference, while the running time of VG depends on the size of the VCF file representing the multiple aligned sequences. This explains the differences in space and time growth between MONI and VG.

Furthermore, VG requires multiple aligned genomes in input, while MONI doesn’t. Hence, MONI would be able to index also non multiple-aligned genomes, e.g., a set of long-read assemblies.

### 4.7 Comparison to Read Aligners: Alignment Analysis

We analyse the results of the alignment of the reads from Mallick et al. (2016) computed in the previous section. In Figure 8 we report, for each length *i* = 1, …, 100, the cumulative number of reads with the longest match of length at least *i* that are aligned. We computed the length of the longest match from the CIGAR string and the MD:Z field of the SAM file. The MD:Z field of the SAM file is not available for VG, hence it was not possible to include it in this analysis. We observe that MONI is the tool having always more reads with a longest match of length at least 26. BWA-MEM has very similar trend with MONI having more reads with a longest matches from 0 to 25. Bowtie2 has a larger gap with respect to MONI and BWA-MEM, for reads with longest matches from 0 to 50. PuffAligner has always fewer longest matches of length at least 40, than all the other tools. There is an evident increase in the number of reads with longer longest matches when moving from `HG` to `HG.10`. By increasing the number of genomes from `HG.10` to `HG.20` in the reference, the curve for MONI and Bowtie2 increases, while for PuffAligner decreases. This trend is preserved increasing even more the number of genomes in the reference. This also demonstrates the importance of indexing a population of genomes, rather than a single genome.

**Fig. 8:**
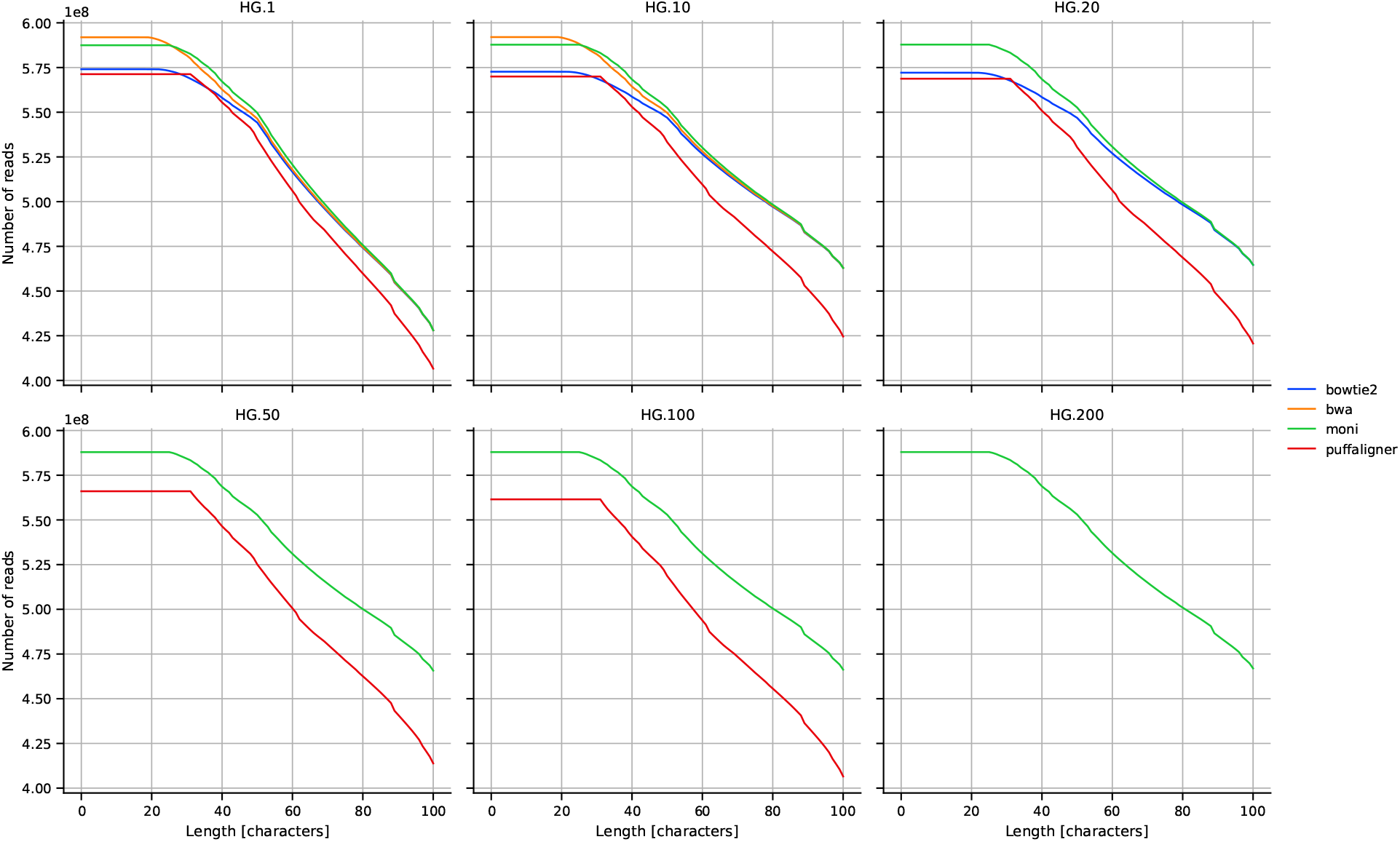
Human genomes dataset cumulative number of reads with the longest match of length at least *i*, for all *i* = 1, …, 100.

## 5 Conclusion

We described MONI, a new indexing method and software tool for finding MEMs between sequencing reads and large collections of reference sequences with minimal memory footprint and index size. While it is not a full-featured aligner – e.g., lacking the ability to compute mapping qualities – MONI represents a major advance in our ability to perform MEM finding against large collections of references. MONI proved to be competitive with graph-based pangenomes indexes as VG. This is promising toward the possibility to perform MEM finding on long-read assemblies. The next step is to thoroughly investigate how to extend these MEMs to full approximate alignments in a manner that is both efficient and accurate.

As explained in this paper, our method hinges on a novel way of computing Bannai et al.’s thresholds with the PFP while simultaneously building the *r*-index. We conclude by noting that there are possible uses of thresholds beyond sequence alignment and these warrant further investigation. For example, as a byproduct of our construction, it is possible to compute the LCP array of the text, which has practical applications in bioinformatics (i.e., SNP finding (Prezza et al., 2019)).

## Acknowledgments

The authors wish to thank Nicola Prezza for the code of rlbwt2lcp.

## Author’s Contributions

MR, TG and CB conceptualized the idea and developed the algorithmic contributions of this work. MR implemented the MONI tool. MR and MO conducted the experiments. MR, MO, CB, and BL assisted and oversaw the experiments and implementation. All authors contributed to the writing of this manuscript.

## Footnotes

4 Formally, given a genome *G*[1..*n*] and read *R*[1..*m*], *R*[*i*..*i* + ℓ – 1] of length ℓ is a MEM of *R* in *G* if *R*[*i..i* + ℓ – 1] occurs in *G* and *R*[*i* – 1..*i* + ℓ – 1] and *R*[*i..i* + ℓ] are not substrings of *G*.

## Bibliography

F. Almodaresi, M. Zakeri, and R. Patro. Puffaligner: An efficient and accurate aligner based on the pufferfish index. Bioinformatics, 2021.

H. Bannai, T. Gagie, and T. I. Refining the r-index. Theoretical Computer Science, 812: 96–108, 2020.

C. Boucher, T. Gagie, A. Kuhnle, et al. Prefix-free parsing for building big BWTs. Algorithms for Molecular Biology, 14(1):13:1–13:15, 2019.

C. Boucher, O. Cvacho, T. Gagie, et al. PFP Compressed Suffix Trees. In 2021 Proceedings of the Symposium on Algorithm Engineering and Experiments (ALENEX), 60–72., 2021.

M. Burrows and D.J. Wheeler. A block sorting lossless data compression algorithm. Technical Report 124, Digital Equipment Corporation, 1994.

A. Dobin, C.A. Davis, F. Schlesinger, et al. STAR: ultrafast universal RNA-seq aligner. Bioinformatics, 29(1):15–21, 2013.

J. Fischer and V. Heun. Space-efficient preprocessing schemes for range minimum queries on static arrays. SIAM Journal on Computing, 40(2):465–492, 2011.

T. Gagie, G. Navarro, and N. Prezza. Fully Functional Suffix Trees and Optimal Text Searching in BWT-Runs Bounded Space. Journal of the ACM, 67(1):2:1–2:54, 2020a.

T. Gagie, T. I, G. Manzini, et al. Rpair: Rescaling RePair with Rsync. In Proceedings of the 26th International Symposium on String Processing and Information Retrieval (SPIRE), 35–44, 2019.

T. Gagie, T. I, G. Manzini, et al. Practical Random Access to SLP-Compressed Texts. In Proceedings of the 27th International Symposium on String Processing and Information Retrieval (SPIRE), 221–231, 2020b.

E. Garrison, J. Siren, A. M. Novak, et al. Variation graph toolkit improves read mapping by representing genetic variation in the reference. Nature Biotechnology, 36(9):875–879, 2018.

S. Gog, T. Beller, A. Moffat, et al. From theory to practice: Plug and play with succinct data structures. In Proceedings of the 13th International Symposium on Experimental Algorithms (SEA), 326–337, 2014.

T. Kasai, G. Lee, H. Arimura, et al. Linear-time longest-common-prefix computation in suffix arrays and its applications. In Proceedings of the 12th Annual Symposium on Combinatorial Pattern Matching (CPM), 181–192, 2001.

A. Kuhnle, T. Mun, C. Boucher, et al. Efficient construction of a complete index for pan-genomics read alignment. Journal of Computational Biology, 27(4):500–513, 2020.

B. Langmead, C. Trapnell, M. Pop, et al. Ultrafast and memory-efficient alignment of short DNA sequences to the human genome. Genome Biology, 10:R25, 2009.

B. Langmead and S. L. Salzberg. Fast gapped-read alignment with Bowtie 2. Nature Methods, 9(4):357–359, 2012.

H. Li. Aligning sequence reads, clone sequences and assembly contigs with BWA-MEM. arXiv, 2013.

H. Li and R. Durbin. Fast and accurate short read alignment with Burrows-Wheeler Transform. Bioinformatics, 25(14):1754–1760, 2009.

H. Li. Minimap2: pairwise alignment for nucleotide sequences. Bioinformatics, 34(18):3094–3100, 2018.

H. Li, X. Feng, and C. Chu. The design and construction of reference pangenome graphs with minigraph. Genome Biology, 21(1):1–19, 2020.

R. Li, H. Zhu, J. Ruan, et al. De novo assembly of human genomes with massively parallel short read sequencing. Genome Research, 20(2):265–272, 2010.

B. Liu, H. Guo, M. Brudno, et al. deBGA: read alignment with de Bruijn graph-based seed and extension. Bioinformatics, 32(21):3224–3232, 2016.

F. A. Louza, S. Gog, and G. P. Telles. Inducing enhanced suffix arrays for string collections. Theoretical Computer Science, 678:22–39, 2017.

A. I. Maarala, O. Arasalo, D. Valenzuela, et al. Scalable Reference Genome Assembly from Compressed Pan-Genome Index with Spark. In Proceedings of the 9th International Conference on Big Data (BIGDATA), 68–84, 2020.

V. Mäkinen and G. Navarro. Rank and select revisited and extended. Theoretical Computer Science, 387(3):332–347, 2007.

V. Mäkinen, D. Belazzougui, F. Cunial, et al. Genome-Scale Algorithm Design: Biological Sequence Analysis in the Era of High-Throughput Sequencing. Cambridge University Press, 2015.

S. Mallick, H. Li, M Lipson, et al. The Simons genome diversity project: 300 genomes from 142 diverse populations. Nature, 538(7624):201–206, 2016.

U. Manber and G. W. Myers. Suffix arrays: a new method for on-line string searches. SIAM Journal on Computing, 22(5):935–948, 1993.

G. Miclotte, M. Heydari, P. Demeester, et al. Jabba: hybrid error correction for long sequencing reads. Algorithms Molecular Biology, 11:10, 2016.

T. Mun, A. Kuhnle, C. Boucher, et al. Matching reads to many genomes with the r-index. Journal of Computational Biology, 27(4):514–518, 2020.

G. Navarro. Compact Data Structures - A Practical Approach. Cambridge University Press, 2016.

G. Nong. Practical linear-time *O* (1)-workspace suffix sorting for constant alphabets. ACM Transactions on Information Systems, 31(3):15, 2013.

N. Prezza and G. Rosone. Space-Efficient Computation of the LCP Array from the Burrows-Wheeler Transform. In Proceedings of the 30th Annual Symposium on Combinatorial Pattern Matching (CPM), 7:1–7:18, 2019.

N. Prezza, N. Pisanti, M. Sciortino, et al. SNPs detection by eBWT positional clustering. Algorithms Molecular Biology, 14(3), 2019.

A. Rhie, S.A. McCarthy, O. Fedrigo et al. Towards complete and error-free genome assemblies of all vertebrate species. Nature, 592(7856):737–746, 2021.

T. F. Smith, M. S. Waterman. Identification of common molecular subsequences. Journal of Molecular Biology, 147(1):195–197, 1981.

E. L. Stevens, R. Timme, E. W. Brown, et al. The public health impact of a publically available, environmental database of microbial genomes. Frontiers in Microbiology, 8:808, 2017.

H. Suzuki and M. Kasahara. Introducing difference recurrence relations for faster semi-global alignment of long sequences. BMC Bioinformatics, 19(1):33–47, 2018.

The 1000 Genomes Project Consortium. A global reference for human genetic variation. Nature, 526:68–74, 2015.

C. Turnbull et al. The 100,000 genomes project: bringing whole genome sequencing to the nhs. British Medical Journal, 361, 2018.

D. Valenzuela and V. Mäkinen. CHIC: a short read aligner for pan-genomic references. bioRxiv, 2017.

D. Valenzuela, T. Norri, N. Välimäaki, et al. Towards pan-genome read alignment to improve variation calling. BMC Genomics, 19(2):123–130, 2018.

M. Vyverman, B. De Baets, V. Fack, et al. A Long Fragment Aligner called ALFALFA. BMC Bioinformatics, 16(1):159, May 2015.

