## Supplement for "MONI: A Pangenomics Index for Finding MEMs"

### MONI: A Pangenomics Index for Finding MEMs Supplementary Material

#### 1 Computation of the BWT given the prefix-free parsing

We provide the pseudocode for the construction algorithm of Kuhnle et al. (2020) which is shown in Algorithm 1. We first provide some additional notation. Given the prefix-free parsing $P$ of $S$ with dictionary $D$, we let $\mathcal{S}$ be the set of the distinct proper phrase suffixes of $S$ of length at least $w$. Given a proper phrase suffix $\alpha \in \mathcal{S}$, we denote by $\mathcal{L}_\alpha$, the set of characters that precede all occurrences of $\alpha$ in $S$. We let $\mathcal{I}_\alpha$ be the set of positions of the phrases whose suffix is $\alpha$ in the $\mathrm{BWT}_P$, and let $\alpha_{First}$ and $\alpha_{Last}$ the first and the last position in $\mathcal{I}_\alpha$. In addition, we denote the number of occurrences of $\alpha$ in $\mathrm{BWT}_P$ as $occs(\alpha)$, i.e. $occs(\alpha) = |\mathcal{I}_\alpha|$. Lastly, given a proper phrase suffix $\alpha$ and an index $i \in \mathcal{I}_\alpha$, we denote by $\alpha_i.\ell = \alpha.\ell$ the length $|\alpha|$ of $\alpha$ and by $\alpha_i.bwt$ as the character preceding $\alpha$ in the phrase. We refer to the character preceding all occurrences of $\alpha$ as $\alpha_i.bwt$ when $|\mathcal{L}_\alpha| = 1$.

**Algorithm 1** Building RL BWT

```
procedure Build RL BWT(P, D)
    i ← 0
    for all α ∈ 𝒮 do
        if |ℒ_α| = 1 then
            i ← Update RL BWT(i, α.bwt, occs(α))
        else
            for all k ∈ ℐ_α do
                i ← Update RL BWT(i, α_k.bwt, 1)
    return RL BWT[1..r]

procedure Update RL BWT(i, a, ℓ)
    if a ≠ RL BWT[i].head then
        i ← i + 1
        RL BWT[i].head ← a
        RL BWT[i].ℓ ← 0
    RL BWT[i].ℓ ← RL BWT[i].ℓ + ℓ
    return i
```

⋆ Both authors should be considered senior authors of the project.

### 2 Computation of the BWT and the thresholds given the prefix-free parsing

In this section we provide the pseudocode for the construction algorithm of Kuhnle et al. (2020) modified to compute in addition the threshold values. The pseudocode is shown in Algorithm 2.

**Algorithm 2** Building RL BWT and *Thresholds*

```
procedure BUILD THRESHOLDS(P, D)
    i ← 0, j ← 1, β ← ε
    for all α ∈ S do
        val ← lcp(α, β)
        UPDATE LCP(val, j)
        if |L_α| = 1 then
            i ← UPDATE RL BWT(i, α.bwt, occs(α))
            UPDATE LCP(val, j)
            j ← j + occs(α)
        else
            prev ← −1
            for all k ∈ I_α do
                if prev ≥ 0 then
                    val ← MIN SLCP(prev, k)
                    val ← val + α.ℓ − w
                UPDATE LCP(val, j)
                i ← UPDATE RL BWT(i, α_k.bwt, 1)
                UPDATE LCP(val, j)
                j ← j + 1
                prev ← k
        β ← α
    return RL BWT[1..r], THR[1..r]

procedure UPDATE LCP(val, pos)
    if val < LCP.val then
        LCP.val ← val
        LCP.pos ← pos

procedure UPDATE RL BWT(i, c, ℓ)
    if c ≠ RL BWT[i].head then
        UPDATE THRESHOLDS(i, c)
        i ← i + 1
        RL BWT[i].head ← c
        RL BWT[i].ℓ ← 0
        LCP ← INIT LCP
    RL BWT[i].ℓ ← RL BWT[i].ℓ + ℓ
    return i

procedure MIN SLCP(a, b)
    if b > a then
        SWAP(a, b)
    return min{SLCP[i] | a < i ≤ b}

procedure UPDATE THRESHOLDS(i, c)
    for all a ∈ Σ \ {RL BWT[i].head} do
        if LCP.val < M[a].val then
            M[a].val ← LCP.val
            M[a].pos ← LCP.pos
    if c is the first c in RL BWT  then
        THR[i].val ← 0
        THR[i].pos ← 0
    else
        THR[i].val ← M[c].val
        THR[i].pos ← M[c].pos
    M[c] ← INIT M
```

### Bibliography

Alan Kuhnle, Taher Mun, Christina Boucher, Travis Gagie, Ben Langmead, and Giovanni Manzini. Efficient construction of a complete index for pan-genomics read alignment. *Journal of Computational Biology*, 27(4):500–513, 2020.
